# Selective inhibition of histone deacetylase 1 and 3 improves motor phenotype and alleviates transcriptional dysregulation in Huntington’s disease mice

**DOI:** 10.1101/2020.10.13.337154

**Authors:** Katharina Hecklau, Susanne Mueller, Stefan Paul Koch, Mustafa Hussain Mehkary, Busra Kilic, Christoph Harms, Philipp Boehm-Sturm, Ferah Yildirim

## Abstract

Huntington’s disease (HD) is an autosomal dominant neurodegenerative disease characterized by a late clinical onset of psychiatric, cognitive, and motor symptoms. Transcriptional dysregulation is an early and central disease mechanism which is accompanied by epigenetic alterations in HD. Previous studies demonstrated that targeting transcriptional changes by inhibition of histone deacetylases (HDACs), especially the class I HDACs, provides therapeutic effects. Yet, their exact mechanisms of action and the features of HD pathology, on which these inhibitors act remain to be elucidated. Here, using transcriptional profiling, we found that selective inhibition of HDAC1 and HDAC3 by RGFP109 repaired the expression of a number of genes, including the transcription factor genes *Neurod2* and *Nr4a2*, and 43% of the gene sets that were dysregulated by mutant Huntingtin expression in the striatum and improved motor skill learning deficit in the R6/1 mouse model of HD. RGFP109-treated R6/1 mice showed improved coordination on the RotaRod over four consecutive trials, while vehicle-treated R6/1 animals displayed no improvement in coordination skills and fell 50 seconds earlier off the rod in the fourth trial. We also found, by volumetric MRI, a widespread brain atrophy in the R6/1 mice at the symptomatic disease stage, on which RGFP109 showed a modest effect. Collectively, our combined work presents new evidence for specific HDAC1 and HDAC3 inhibition as a therapeutic strategy for alleviating the phenotypic and molecular features of HD.

## Introduction

Huntington’s disease (HD), an autosomal dominant neurodegenerative disease caused by CAG repeat expansions in the exon I of the *Huntingtin* (*HTT*) gene (The Huntington’s Disease Collaborative Research Group, 1993), is characterized by progressive psychiatric, motor, and cognitive symptoms and is fatal. In *HTT* exon 1, normal individuals have 7-34 CAG repeats, while HD patients display more than 40 and in juvenile cases even more than 100 CAG repeats (Duyao et al., 1993). Currently, no disease-modifying treatment is available. Mutant *HTT* gene causes brain region-specific neuronal dysfunction and degeneration that is most prominent in the striatum during early disease stages and spreads to other brain regions when the disease progresses (Tabrizi et al., 2012).

Transcriptional dysregulation is an early and central pathogenic mechanism in Huntington’s disease which has been demonstrated in cell and animal models of HD as well as in human HD brain (Cha, 2000; Hodges et al., 2006; Seredenina and Luthi-Carter, 2012). Transcriptional repression of many genes coding for neurotransmitters, neurotrophins, and their receptors is a hallmark of HD, while genes that are part of stress-response pathways were shown to be upregulated in HD. Some of the key neuronal genes that are consistently reported to be repressed across HD patients and animal models include brain-derived neurotrophic factor (*Bdnf*), preproenkephalin (*Penk*), dopamine receptor 2 (*Drd2*), and dopamine receptor 1a (*Drd1*). Accordingly, it has been shown that overexpression of *Bdnf* and *Penk* is neuroprotective and improves disease outcome in HD (Bissonnette et al., 2013; Zuccato et al., 2005; Zuccato et al., 2011). Furthermore, dopamine agonist and antagonists play an important role in symptomatic treatment of HD patients (Cepeda et al., 2014; Coppen and Roos, 2017; Rangel-Barajas et al., 2015). However, the mechanisms driving transcriptional dysregulation in HD are not fully understood. It was suggested that mutant HTT directly interacts with transcription factors and DNA (Benn et al., 2008). On the other hand there is strong evidence for epigenetic mechanisms such as histone acetylation, histone ubiquitination, histone trimethylation (H3K9me3, H3K4me3), and DNA methylation contributing to selective changes in gene expression in HD pathogenesis (Bett et al., 2009; Ferrante et al., 2004; Hervas-Corpion et al., 2018; Ryu et al., 2006; Seo et al., 2008; Vashishtha et al., 2013).

Histone hyperacetylation by histone acetyltransferases (HATs) is generally associated with gene expression and histone hypoacetylation by histone deacetylases (HDACs) with gene repression (Kurdistani et al., 2004). Global as well as gene specific histone hypoacetylation at promoters of down-regulated genes were shown in various HD models (Sadri-Vakili et al., 2007; Yildirim et al., 2019). Mutant HTT leads to hypoacetylation by directly binding HATs such as CBP, reducing their activity and resulting in gene repression (Cong et al., 2005; Jiang et al., 2006; Nucifora et al., 2001; Steffan et al., 2000). Overexpressing CBP reverses these effects and leads to decreased mutant HTT toxicity (Jiang et al., 2006; Steffan et al., 2000). Similar neuroprotective effects are achieved by inhibiting the opponent, HDACs. HDAC inhibition has been shown to ameliorate transcriptional changes in HD and improve behavioral deficits across different experimental models (Butler and Bates, 2006; Ferrante et al., 2003; Ferrante et al., 2004; Hockly et al., 2003; Kazantsev and Thompson, 2008; Pallos et al., 2008; Ryu et al., 2006; Steffan et al., 2001). Beyond HD, such chromatin targeting strategies exert therapeutic effects in other neurological conditions such as ischemic injury, Alzheimer disease, and Amyotrophic lateral sclerosis as previously reported by us and others (Faraco et al., 2006; Harrison and Dexter, 2013; Meisel et al., 2006; Ryu et al., 2005; Schweizer et al., 2015; Yang et al., 2017; Yildirim et al., 2008). These findings provide significant support for the disease-modifying therapeutic potential of epigenome-targeting strategies such as inhibition of histone deacetylation.

Initially, HDAC inhibitors have been studied and applied in cancer research identifying non-selective HDAC inhibitors such as Trichostatin A (TSA) and Vorinostat as anti-cancer drugs. In recent years, however, selective HDAC inhibitors have been developed (Wagner et al., 2015) and there is growing evidence that especially HDAC class I inhibitors are effective in suppressing pathogenic mechanisms in mouse models of HD (Chen et al., 2013; Chiu et al., 2011; Jia et al., 2012; Jia et al., 2016; Lim et al., 2011; Suelves et al., 2017; Thomas et al., 2008). RGFP109 is a HDAC class I inhibitor selective for HDAC1 and HDAC3. RGFP109 confirmed safety in a recent Phase I clinical trial in patients of Friedreich Ataxia Syndrome and enhanced the mRNA levels of the key disease gene frataxin (FXN) in patient blood mononuclear cells indicating its potentials for disease amelioration (Soragni et al., 2014). In another study, RGFP109 treatment led to significantly decreased levels of L-dopa induced dyskinesia in a Parkinson’s disease marmosets model (Johnston et al., 2013). A short-term treatment of HD R6/2 mice with RGFP109 modified the expression levels of 4 out of 13 measured disease-associated genes in the striatum (Jia et al., 2012). So far, a comprehensive investigation of the effects of RGFP109 on different aspects of HD, such as behavioral disease phenotypes or brain atrophy, has not been carried out.

In the present study, we tested the therapeutic effects of the selective HDAC inhibitor RGFP109 in the R6/1 mouse model of HD. The outcome of HDAC1 and HDAC3 inhibition was assessed by a set of behavioral tests and genome-wide transcriptional analysis of the striatum. Moreover, we performed volumetric magnetic resonance imaging (MRI) of the brain in living animals. Our results demonstrate that inhibition of HDAC1 and HDAC3 alleviates primarily the short-term motor skill learning deficits, accompanied by a partial repair effect on global gene expression changes in the striatum of R6/1 mice. MR imaging showed in addition, that RGFP109 treatment exerted modest changes on the atrophy of specific brain regions in the R6/1 mice. Collectively, these findings present evidence for beneficial effects of specific HDAC1 and HDAC3 inhibition on several aspects of disease pathology in HD mice.

## Results

### HDAC1 and HDAC3 inhibition by RGFP109 ameliorates motor learning deficits typical of HD in R6/1 mice

R6/1 transgenic mice faithfully recapitulate many of the disease features of human HD, such as transcriptional dysregulation, progressive impairments of both motor and cognitive functions (Brooks et al., 2012; Hodges et al., 2008; Naver et al., 2003; Yildirim et al., 2019), brain atrophy and mutant HTT accumulation (Bayram-Weston et al., 2012). In our study, to evaluate if the specific HDAC inhibitor RGFP109 could alleviate the transcriptional dysregulation in HD *in vivo* and improve the characteristic neuroanatomical HD features and behavioral deficits, we treated R6/1 mice (11 to 14 weeks of age) with 30 mg/kg (i.p.) RGFP109 five times a week for three weeks (total of 23 days) (Figure 1A). R6/1 mice display a range of characteristic behavioral deficits such as motor abnormalities, learning and memory impairments, and reduced level of anxiety (Brooks et al., 2012; Hodges et al., 2008; Naver et al., 2003). Thus, we examined the R6/1 mice using different behavioral tests including RotaRod, open field, and elevated plus maze during the course of the study (Figure 1A).

**Figure 1.**
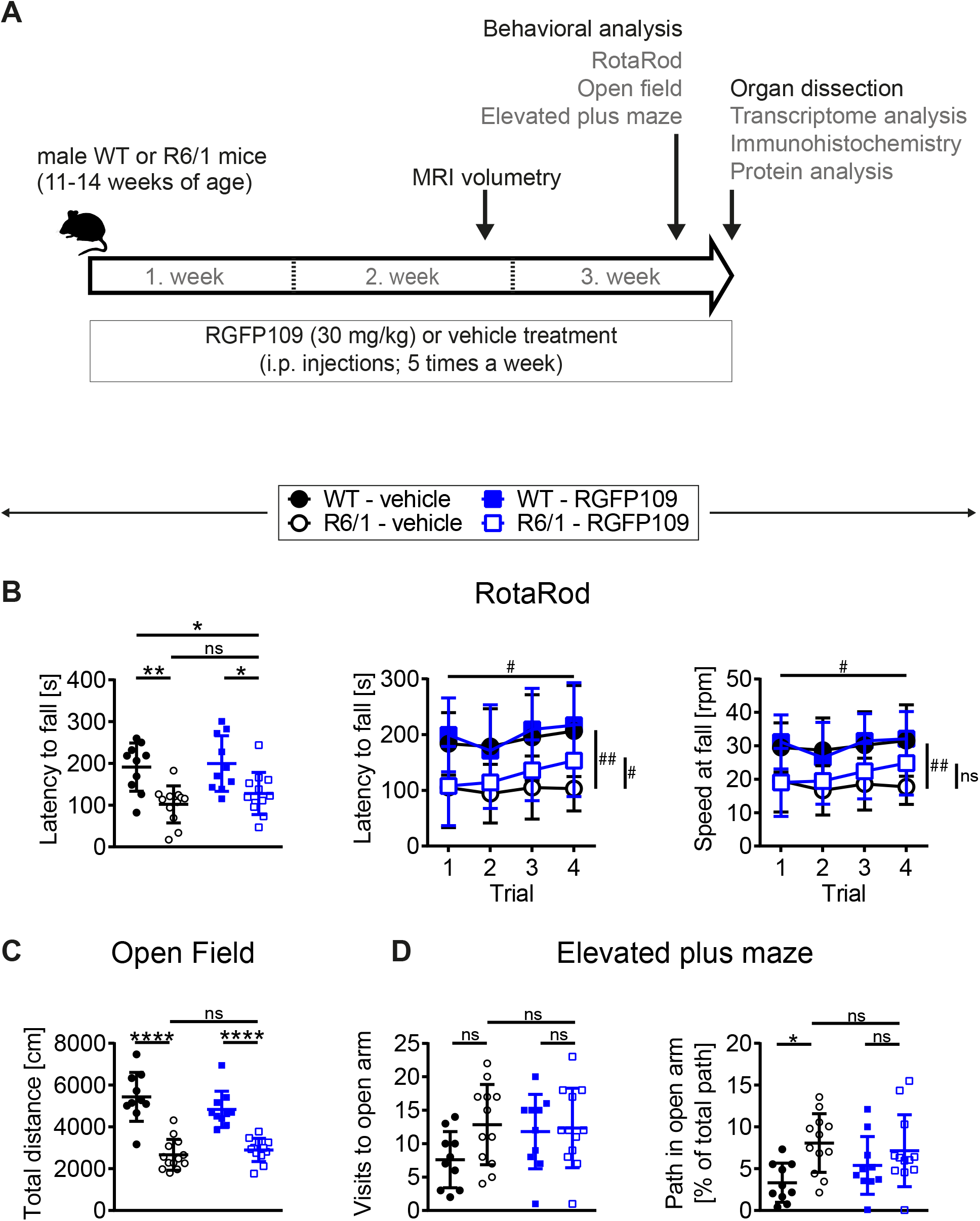
RGFP109 treatment improves motor learning in R6/1 mice. (A) Schematic of treatment and experimental procedure. (B) Motor performance and learning on a RotaRod: Latency to fall as mean of all trials, latency to fall and speed at fall as learning curve over the course of four subsequent trials. (C) Open field exploration evaluating general locomotor activity measured as total distance travelled. (D) Anxiety-related behavior determined by elevated plus maze test showing visits to open arm and path travelled in open arm normalized to the total path. WT – wild type. Data represent mean+SD; n=10 [WT-vehicle], n=12(13, Open Field) [R6/1-vehicle], n=10 [WT-RGFP109], n=12 [R6/1-RGF109]; *p ≤ 0.05, **p ≤ 0.01, ***p ≤ 0.001, ****p ≤ 0.0001, ns not significant by two-way ANOVA with Tukey’s multiple comparisons test (B-left graph, C, and D); ^#^q ≤ 0.05, ^##^q ≤ 0.01, ns not significant by mixed-effects analysis with two-stage linear step-up procedure of Benjamini, Krieger and Yekutieli as post-tests by controlling the False Discovery Rate (individually comparing each group to each other group) (B-learning curves: comparison of trial 4 between groups - indicated vertically, comparison of trial 4 *vs*. trial 1 for R6/1-RGFP109 – indicated horizontally).

For evaluating the effect of HDAC1 and HDAC3 inhibition on motor coordination, balance, and motor skill learning in R6/1 mice, we performed the accelerating RotaRod test. Figure 1B (left) depicts the latency to fall as mean of four consecutive trials for each animal. Comparison of the vehicle-treated R6/1 and wild type animals revealed an HD-characteristic phenotype with significantly shorter time to stay on the rod in R6/1s. The latency to fall was significantly reduced by the main genotype effect (F_1,40_=23.6 with p<0.0001) while treatment had only a minor effect on motor coordination (F_1,40_=1.09 with p=0.304). However, mice with treatment stayed 26 s longer on the rod, albeit not being significant (mean of all trial: 101.8±44.2 s for R6/1-vehicle and 127.8±50.5 s for R6/1-RGFP109; p=0.651) (Figure 1B, left). This was also observed for the speed at fall (mean of all trials: 18.1±5.7 rpm for R6/1-vehicle and 21.4±7.4 rpm for R6/1-RGFP109; p=0.672) (Figure S1A). In contrast, inhibition of HDAC1 and HDAC3 in R6/1 mice led to a significant improvement in RotaRod performance over the course of the four consecutive trials, indicating enhanced motor learning skills in HDACi-treated R6/1 animals (q=0.017 for trial 4 *vs*. trial 1 for latency to fall; q=0.026 for trial 4 *vs*. trial 1 for speed at fall) (Figure 1B, middle and right). Notably, RGFP109 treatment in R6/1 animals resulted in a significant increase in time to stay on the rod compared to vehicle-treated R6/1s in the fourth trial (103.1±40.0 s for R6/1-vehicle and 153.3±64.6 s for R6/1-RGFP109 and a difference of 50.2 s; q=0.048) (Figure 1B, middle). Albeit not statistically significant, a similar trend was observed for the speed at fall (mean difference=7.08; q=0.052) (Figure 1B, right). Vehicle-treated R6/1 mice as well as vehicle- and RGFP109-treated wild type animals did not exhibit significant changes in motor learning abilities during the four trials. These findings indicate that HDAC1 and HDAC3 inhibition by RGFP109 ameliorates motor learning deficits in R6/1 mice.

We next analyzed if the general locomotor activity in the open field was also affected by RGFP109 treatment in R6/1 mice. Locomotor activity is known to be impaired in the R6/1 line as well as in other HD models (Hodges et al., 2008; Naver et al., 2003). Accordingly, we observed a highly significant reduction in the total distance travelled as well as in the average velocity and vertical activity (supported rearing) in vehicle-treated R6/1 animals compared to vehicle-treated wild type littermates indicating a hypoactive phenotype in HD animals (Figure 1C, Figure S1B). However, no significant differences were found between vehicle- and RGFP109-treated R6/1 mice for any of these parameters. Next, we performed the elevated plus maze test, which has been developed for measuring anxiety in rodents (Walf and Frye, 2007). As previously described (Naver et al., 2003), R6/1 mice showed trends to visit the open arms more frequently (7.6±4.2 for WT-vehicle, 12.8±6.0 for R6/1-vehicle; p=0.137), spend more time (13.6±11.2 sec for WT-vehicle, 26.3±13.6 sec for R6/1-vehicle; p=0.163), and travel longer distances (38.8±29.2 cm for WT-vehicle, 78.0±45.5 cm for R6/1-vehicle; p=0.198) in the open arms regardless of treatment. Taking the reduced locomotor activity of R6/1s into account, vehicle-treated R6/1 mice displayed a significantly less anxious phenotype than vehicle-treated wild type mice (visits, time, and path in open arm normalized to total path). Treatment with RGFP109 did not exert a significant effect on this phenotype neither in R6/1 nor in wild type mice (Figure 1D, Figure S1C).

Lastly, abnormal limb clasping upon tail suspension can be used to demonstrate the presence of a neurological phenotype in HD mice. At the end of our study, before the sacrifice of mice (at 14-17 weeks of age), several R6/1 animals exhibited first degree forelimb clasping (score 0.5). However, RGFP109 administration did not affect the clasping behavior (Figure S1D). During the course of the study, body weight of the animals was not influenced by HDAC inhibitor administration indicating that the treatment was in general well tolerated. As expected, a reduction in the weight of the R6/1 mice was observed with disease progression, on which HDACi treatment showed no effect (Mangiarini et al., 1996; Naver et al., 2003) (Figure S1E).

Altogether, these results demonstrate that RGFP109 treatment alleviates primarily the motor skill learning deficits while not affecting the general locomotor activity or anxiety-related phenotypes in HD mice.

### RGFP109 treatment alleviates, in part, the global gene expression changes in the striatum of R6/1 mice

Aberrant transcriptional regulation in brain, especially in striatum and cortex, is an early and central feature of HD pathogenesis. Among the genes with aberrant expression in HD striatum are transcriptional regulators (e.g. *Fos, Egr1, Npas4*, and *Polr2a*) as well as key neuronal genes important for neurotransmitter signaling (e.g. *Drd1, Drd2, Grin3a*, and *Ppp1r1b*) and synaptic plasticity (e.g. *Arc* and *Syp*) (Seredenina and Luthi-Carter, 2012; Vashishtha et al., 2013; Yildirim et al., 2019). To examine the impact of RGFP109 treatment on transcriptional dysregulation, we conducted transcriptome analysis of the striatum by RNA-sequencing (RNA-seq) (n=5 for WT-vehicle, n=6 for R6/1-vehicle, n=6 for WT-RGFP109, n=7 for R6/1-RGFP109). All animals were sacrificed 18 h after the final injection and the striatum was used for RNA isolation and subsequent analyses (Figure 1A).

Analysis of the RNA-seq data revealed that 1461 genes were significant differentially expressed in the R6/1 striatum at the age of sacrifice compared to wild type mice in the vehicle-treated group (FDR, q<0.1; log2 FC > |0.5|) with 1171 genes (80%) being downregulated in R6/1 animals (Dataset S01). As expected, among these, key HD genes such as *Drd1, Drd2, Ppp1r1b, Penk*, and *Adora2a* were downregulated. Analysis of the RNA-seq data from the RGFP109-treated R6/1 striata showed that the HDACi treatment significantly changed the expression of 43 genes in R6/1 animals (FDR, q<0.1; log2 FC > |0.5|), of which 36 genes were also differentially expressed in R6/1s compared to wild types (Dataset S01). 14 out of these 36 genes were changed towards wild type expression levels upon RGFP109 treatment in R6/1 animals (Figure 2A, highlighted in red). Among these are transcription factor genes such as *Neurod2, Neurod6*, and *Nr4a2*, the transcriptional regulator *Satb2*, the thyrotropin-releasing hormone *Trh*, the neuropeptide-like molecule *Nxph3*, and the nicotinic acetylcholine receptor *Chrna4* genes. We validated the RNA-seq findings by quantitative RT-PCR for several typical HD genes (*Drd1, Drd2, Ppp1r1b, Adora2a, Arc, Egr1, Polr2a*, and *Grin3a*), for the effect of HDAC1 and HDAC3 inhibition on the transcriptional changes in R6/1s (*Nr4a2, Satb2, Folr1*, and *Otx2*) (although not significant in qRT-PCR data), as well as for several genes that were previously connected to motor skill learning behavior in rodents (*Dpysl2, Wars*, and *Cpne5*) (D’Amours et al., 2011) in the striatum of vehicle- and RGFP109-treated wild type and R6/1 mice (Figure S2A). The complete lists of significant differentially expressed genes for the R6/1 and wild type comparison as well as for the treatment effect in R6/1 mice in RNA-seq data are presented in Dataset S01.

**Figure 2.**
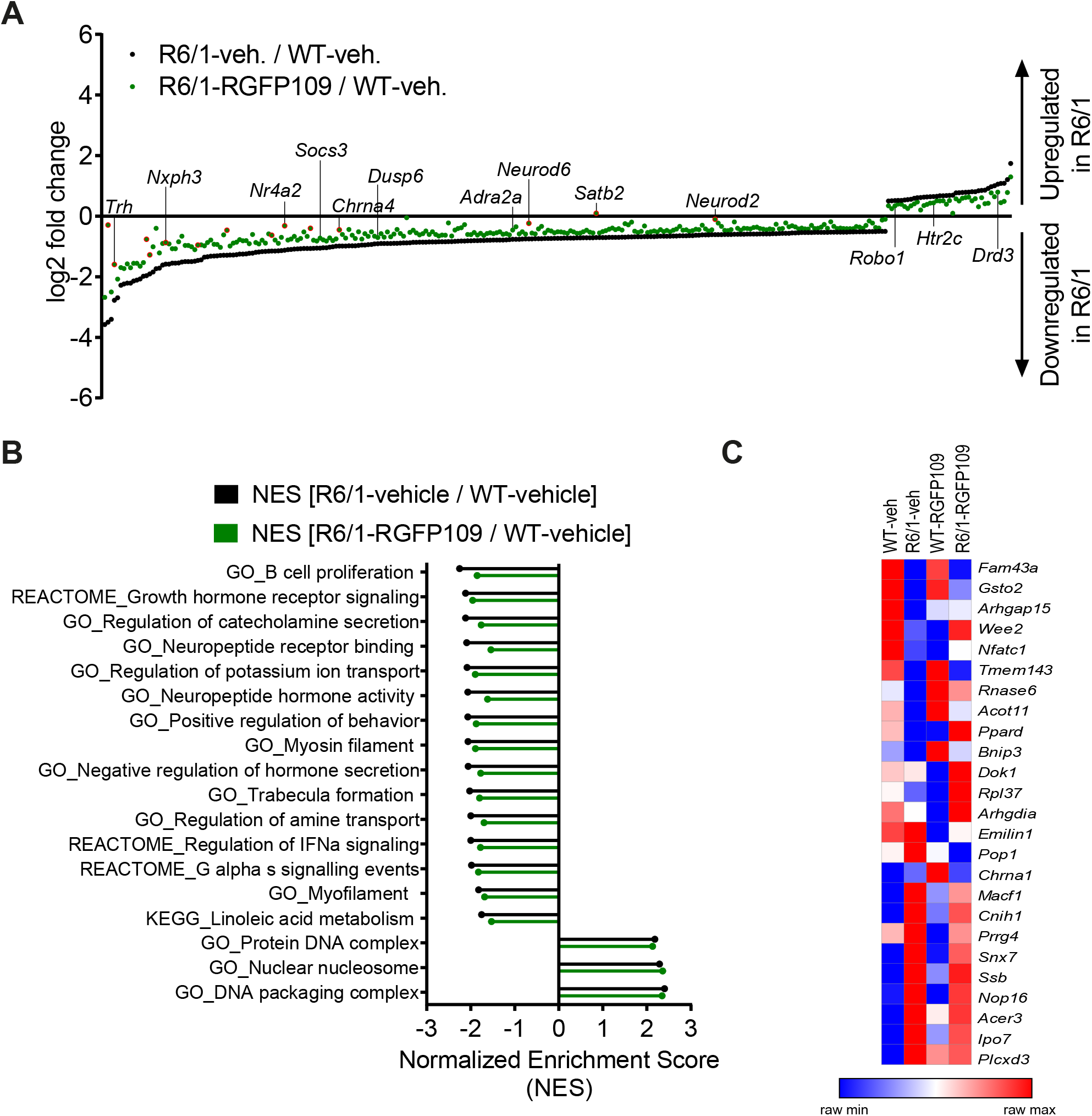
Inhibition of HDAC1 and HDAC3 by RGFP109 has a partial repair effect on global gene expression changes in the striatum of R6/1 mice. (A) 283 genes, significant differentially expressed between WT and R6/1 mice (black) (*Cuffdiff 2*; FDR, q<0.1; log2 FC >|0.5|), are changed towards wild type expression levels in RGFP109-treated R6/1 animals by at least 20% using no cutoff criteria on the data for statistical significance (green). Significant differentially expressed genes upon RGFP109 treatment in R6/1 mice (R6/1-RGFP109 / R6/1-vehicle; FDR, q<0.1; log2 FC >|0.5|) are framed in red. Selected genes are labeled. (B) Gene set enrichment analysis (GSEA - Broad Institute) for gene ontology, KEGG pathway, and REACTOME pathway gene sets. Genes were ranked based on log2 fold changes. The bar graph shows gene sets with significant different normalized enrichment scores (NES) between WT-vehicle and R6/1-vehicle mice (FDR, q<0.1). NESs of RGFP109-treated R6/1 compared to WT-vehicle mice are plotted for the same gene sets. Gene sets with positive NES and the top 15 gene sets with highest difference between both comparisons for negative NES are plotted. (C) Heatmap for gene expression levels of 25 genes, that show a change towards wild type levels by RGFP109 treatment in R6/1 mice, from a list of genes whose expression were restored by HDACi treatment in a previous study (Thomas et al., 2008). Mean FPKM value/group is shown. Each raw is a gene and each column is a group. n=5 [WT-vehicle], n=6 [R6/1-vehicle], n=7 [R6/1-RGFP109].

Further analysis of the datasets, without imposing cutoff criteria on the data for statistical significance, revealed a trend for a larger potential effect of RGFP109 treatment on expression of the dysregulated genes in R6/1 striatum, as illustrated in Figure 2A. Of the 1171 downregulated genes in the vehicle-treated R6/1s, expression of 21% (244) were enhanced by >20% by the HDACi treatment of R6/1s and among these were genes such as adrenoceptor alpha 2a (*Adra2a*) and 1b (*Adra1b*), suppressor of cytokine signaling 3 (Socs3), and dual specificity phosphatase 6 (*Dusp6*), whose expression showed a shift towards wild type levels. Similarly, expression of 13% (39) of the genes upregulated in the vehicle-treated R6/1 striatum were reduced by >20% by the RGFP109 treatment and some of these genes were 5-hydroxytryptamine (serotonin) receptor 2c (*Htr2c*), roundabout guidance receptor 1 (*Robo1*), and dopamine receptor d3 (*Drd3*) (Figure 2A and Figure S2B). To get a better functional view of genes that were alleviated by HDACi treatment in R6/1 animals, we performed gene ontology (GO) enrichment analysis of these genes that were changed towards wild type expression levels in RGFP109-treated R6/1 animals by at least 20%. These genes were enriched for several GO terms that are relevant for brain physiological processes, such as neuropeptide hormone activity, regulation of secretion, and second-messenger-mediated signaling (Figure S2C). Dataset S02 shows the complete list of enriched GO terms (FDR, q<0.1).

Next, we examined if expression levels of certain sets of genes associated with specific functions, phenotypes, or pathways would be more affected by HDAC1 and HDAC3 inhibition than individual genes. Using log2 fold changes for ranking genes, we performed gene set enrichment analysis (GSEA) to find the gene sets associated to the functional transcriptional changes in HD and those induced by HDAC1 and HDAC3 inhibition. Indeed, normalized enrichment scores of 43% (24 out of 55) of the gene sets that were negatively enriched in disease (R6/1-vehicle / WT-vehicle comparison) were changed by at least 5% towards wild type levels in RGFP109-treated R6/1 mice (enrichment scores shifting towards 0 comparing drug-treated R6/1s *vs*. vehicle-treated wild type mice). Among these were gene sets associated with neuropeptide receptor binding, negative regulation of hormone secretion, and regulation of IFNα signaling, that were increased towards wild type levels. In contrast, among the positively enriched gene sets in R6/1, only protein dna complex and dna packaging complex were lowered towards wild type levels upon treatment (by 2.4% and 2.3%, respectively). Of note, the neuroactive ligand receptor interaction gene set showed further enhancement of the disease-associated enrichment pattern upon HDAC1 and HDAC3 inhibition in R6/1 animals (by 6.4%), while the remaining gene sets enriched in disease (31) were not changed or changed by less than 5% by RGFP109 treatment and these include actin filament based movement and potassium channel complex gene sets (Figure 2B, Dataset S02).

We next compared our results with a previous report which examined the effects of a similar HDACi, HDACi 4b, on genome-wide gene expression profiles in the striatum of R6/2 mice using microarray (Thomas et al., 2008). Of 56 genes, which were shown to be restored in the striatum by HDACi 4b treatment of R6/2 mice, 42 were detected by RNA-seq in our study. Of these 42 genes, only 6 were significantly dysregulated in the striatum of R6/1 mice at the age of 14-17 weeks, which corresponds to a less progressed disease stage compared to the R6/2s, and 2 of these genes, *Arhgap15* and *Gsto2*, were significantly restored by RGFP109 treatment of the R6/1s in our study. To explore a more inclusive list of potentially therapeutically relevant genes, we examined all expression changes, regardless of their statistical significance, which revealed that of the 42 genes previously reported to be restored by HDACi 4b, 25 showed a change toward the WT levels after RGFP109 administration in R6/1 mice in our study (Figure 2C, Dataset S01), providing a list of genes whose expression levels are consistently ameliorated by specific HDAC1/3 inhibitors across different HD models.

Collectively, RGFP109 treatment at least to a certain extent, repairs the transcriptional effects of mutant Huntingtin gene expression, causing a significant change in the expression of 43 genes, in the striatum of R6/1 mice and induced collective changes in the expression of gene sets, some of which were associated to biological functions and processes that are relevant for neuronal physiology and HD pathogenesis.

### Effects of RGFP109 treatment on the HD-associated atrophy of specific brain areas in the R6/1 mice measured by volumetric MRI

One hallmark of HD is neuronal degeneration which manifests predominantly in the striatum, particularly the caudoputamen and dorsal striatum in HD patients. In addition, brain atrophy of variable severity can be observed in several other brain regions, such as the cerebral cortex, total white matter, amygdala, hippocampus, and brainstem (Rosas et al., 2003; Tabrizi et al., 2012). The R6/1 mouse model closely recapitulates the neuronal degeneration seen in HD patient brain. These mice exhibit progressive brain atrophy in the striatum (especially the posterior striatum) and cortex (especially in the retrosplenial areas) as well as a subtle expansion of posterior ventricular spaces (Rattray et al., 2013).

By registering the Allen brain atlas to T2w MR images (Koch et al., 2019) we used an unbiased approach to detect regional and sub-regional changes in R6/1 mice upon RGFP109 treatment (Figure 1A). This analysis revealed 419 brain regions and sub-regions (55% of all analyzed regions) that showed a significant volume difference between vehicle-treated R6/1 and wild type mice at the age of 13-16 weeks of age (FDR, q<0.1) (Figure 3A, Figure S3A). Of these, the majority (403 regions, 96%) exhibited a significant volume decrease in R6/1 mice including striatum, pallidum, hippocampal region, and piriform area. In contrast, 16 regions (4%) showed a significant volume increase in HD mice, e.g. copula pyramidis, orbital area (medial part, layer 1 and 2), and supratrigeminal nucleus (Figure S3E and Dataset S03). RGFP109 treatment did not induce any significant changes in the volumes of the afore-mentioned 419 brain substructures in R6/1 mice at this age. However, 61 regions (15%) with significant decrease or increase in R6/1 animals showed non-significant trends towards a reversal of the volume change upon drug treatment by at least 20% (Figure 3A, framed red). Some of these regions include central amygdalar nucleus (medial part), pallidum (ventral region), and prelimbic area. Dataset S03 contains the complete list of brain regions with significant volume difference in R6/1 animals and those that are impacted by the treatment.

**Figure 3.**
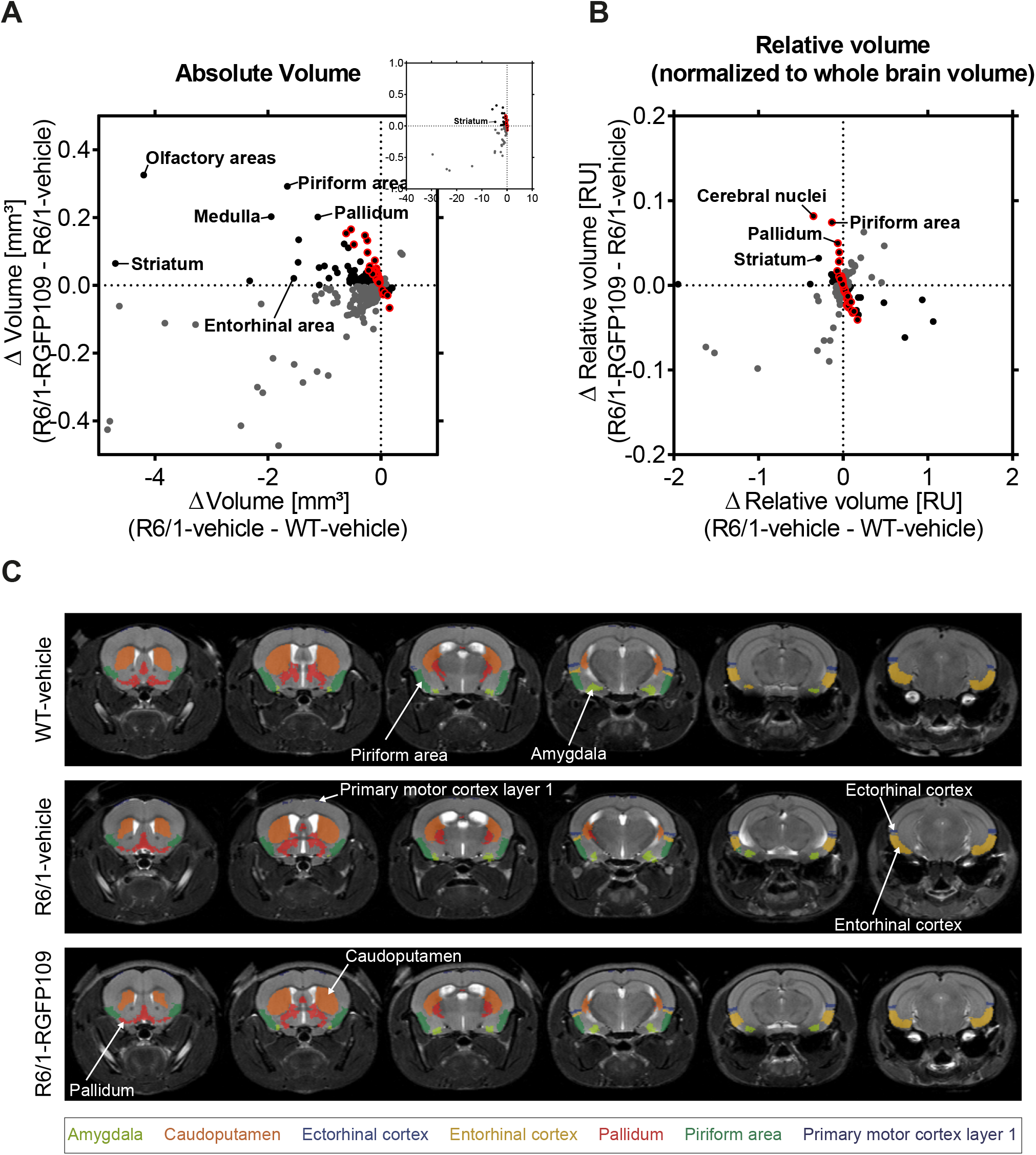
Effect of RGFP109 treatment on HD-associated atrophic brain substructures in R6/1 mice. Mouse brain MR images were registered to the Allen brain atlas. Volumes of substructures (absolute or normalized to whole brain volumes) were analyzed by two-sample t-test (FDR, q<0.1). (A) Mean absolute volume differences of brain regions with significant volume change in R6/1 compared to WT mice (419). (B) Mean relative volume differences of brain regions with significant volume change in R6/1 compared to WT mice (220). (A, B) Black: substructures affected positively by drug (not significant; red frame: >20% of R6/1-WT volume difference); Gray: substructures affected negatively by drug (not significant). (C) MR images, exemplarity shown for three individuals; selected regions are color-coded. Anatomical labels are based on the Allen brain atlas.

As the overall brain volumes in R6/1 mice were reduced (Figure S3E, first graph), we utilized a second approach that corrects the regional and sub-regional volumes by the whole brain volume. This resulted in 220 regions (29% of all analyzed regions) showing a significant change in R6/1 animals (Figure 3B, Figure S3B). 144 brain regions with significant volume change were shared between the two analysis strategies (Figure S3C). Similar to the absolute volume analysis, RGF109 treatment did not show any significant effects on volumes of brain regions in R6/1 mice after correcting for the whole brain atrophy. Non-significant trends towards an improved volume change of at least 20% upon drug treatment in R6/1 mice was detectable for 25% (55/220) of the regions showing a significant volume difference in R6/1 mice (Figure 3B, framed red). 17 brain regions and sub-regions positively affected by drug treatment by more than 20% in R6/1s were shared in both analysis strategies. These include piriform area (molecular layer), ectorhinal area, intermediodorsal nucleus of the thalamus, and prelimbic area (layer 5) (Figure S3D, Figure S3E, Dataset S03). Figure 3C depicts selected regions that were previously implicated in motor skill learning (amygdala, caudoputamen, ectorhinal cortex, entorhinal cortex, pallidum, piriform area, primary motor cortex) in representative examples of wild type, R6/1 and HDACi-treated R6/1 mice (Badea et al., 2019; Scholz et al., 2015; Tamakoshi et al., 2014).

In summary, matching the Allen brain atlas to MR images allowed us to investigate the HD mouse neuroanatomy and the impact of RGFP109 treatment on it in great detail. While we detected a large number of brain regions and sub-regions with significant volume change in R6/1 animals, HDAC1 and HDAC3 inhibition exhibited only trends towards slowing these volumetric differences in the brains of R6/1 mice at this progressed disease stage.

### RGFP109 treatment does not affect aggregate formation or bulk histone H3K27acetylation levels in R6/1 mice

HD is characterized by mutant HTT protein aggregate formation, which displays a histopathological basis of transcriptional dysregulation, neuronal degeneration, and behavioral deficits. Given the effect of RGFP109 on short-term motor skill learning, we examined whether RGFP109 treatment has an impact on this neuropathological hallmark of HD. For this purpose, we analyzed EM48 immunoreactivity in striatal sections visualizing aggregated mutant HTT. We analyzed 8 separate images per section, each covering a 34192 μm^2^ area with 123 DAPI+ cells in average in the dorsal stratum directly lateral to the lateral ventricle. By using ImageJ software, we defined particles larger than 15 μm^2^ as nuclei. Blinded manual counting of the number of aggregates showed that RGFP109 did not affect the number of mutant HTT protein aggregates in R6/1 mice (Figure S4A).

To examine the effects of RGFP109 treatment on global histone acetylation, we determined bulk acetylation patterns at lysine 27 of histone H3 (H3K27ac) in protein extracts of striatal tissue *via* immunoblotting. Increased H3K27ac levels can be found at active promoters and enhancers indicating high transcriptional activity (Wang et al., 2008). Our Western blotting analysis showed no differences in acetylated H3K27 levels between R6/1 and wild type mice, regardless of treatment. As control for equal histone amounts, we determined total H3 levels from the same samples. Similarly, we did not observe differences between the groups (Figure S4B). While we did not detect increases in global H3K27ac patterns in striatum by the HDACi treatment, temporally dynamic and site-specific changes in acetylation levels not detected by immunoblotting of total tissue could be associated to the transcriptional changes observed in HD.

## Discussion

This study shows that treating HD mice with a specific HDAC1 and HDAC3 inhibitor has beneficial effects on multiple aspects of HD. Inhibiting histone modifying enzymes, in particular class I HDACs, has been previously shown to improve disease features, such as the phenotypic and neuropathological deficits and dysregulation of selected genes, in various models of HD (Chen et al., 2013; Chiu et al., 2011; Jia et al., 2012; Jia et al., 2016; Lim et al., 2011; Suelves et al., 2017; Thomas et al., 2008). The goal of our study was to evaluate the effects of the specific HDAC1 and 3 inhibitor RGFP109 in an unbiased approach on the genome-wide transcriptional dysregulation and on the phenotypic deficits in HD mice using the R6/1 model. Further, by performing MRI measurements on living mice, we conducted a detailed analysis of the brain volumetric changes in the R6/1 mice. Our findings demonstrate that the specific inhibition of HDAC1 and HDAC3 by RGFP109 improves primarily the motor learning skills and alleviates, at least to a certain extent, the dysregulation of a number of genes and gene sets in the R6/1 mice. By volumetric MRI, we detected extensive atrophic changes in the brains of the R6/1 mice, on which RGFP109 exerted a statistically non-significant modest effect. Given our results and data from others, early targeting of transcriptional dysregulation by specific HDAC inhibition may alleviate key aspects of HD pathology.

In a previous study, a short-term treatment regime with RGFP109 (subcutaneous injections of 150 mg/kg/day for three days) partially reversed four selected disease-associated genes in the striatum in HD R6/2 mice as measured by PCR (Jia et al., 2012). However, the effects of RGFP109 treatment on the behavioral deficits and on gene expression throughout the genome as well as on the brain atrophy have not been evaluated in HD. Here, using the R6/1 mice, which is less progressive than the R6/2 model, we chose to use a lower therapeutic dose of 30 mg/kg based on a previous Parkinson’s disease study (Johnston et al., 2013) and applied a longer treatment regime (three weeks with five injections per week), aiming to minimize potential toxic side effects by reducing high drug exposure.

Among the most characteristic behavioral deficits in HD mice are motor impairment and deficit in motor skill learning, which are caused by the dysfunction of corticostriatal circuits (Backman et al., 1997; Cybulska-Klosowicz et al., 2004; Lawrence et al., 2000; Mazarakis et al., 2005). The RotaRod task is a useful marker for detecting early HD phenotypes in R6/1 animals from as early as 8 weeks of age (Brooks et al., 2012). Our study demonstrates that 3-week-long treatment with 30 mg/kg RGFP109 ameliorates motor skill learning deficits in this mouse model of HD. While we obtained this finding by testing the animals on the RotaRod for up to four trials on the same day, in future studies, it would be important to analyze the animals across several days to assess learning over longer time frames as well as to use modified RotaRod protocols or other behavioral tests to better distinguish motor skill learning from motor coordination and balance. In line with our findings, Suelves *et al*. demonstrated that selective HDAC3 inhibition by RGFP966 in the Hdh^Q7/Q111^ knock-in mouse model of HD (50 mg/kg of RGFP966 three times per week from 3 to 6.5 months) prevents corticostriatal-dependent motor learning deficits from trial four on testing animals for four times per day for three consecutive days (Suelves et al., 2017). Similarly, treatment of N171-82Q HD mice with RGFP966 over a period of 10 weeks with 10 or 25 mg/kg per week showed improved motor function, accompanied by neuroprotective effects on striatal volume, and significant alterations of the expression of 3 immune pathway genes (chemokine (C-C motif) ligand 17 (*Ccl17*), macrophage migration inhibitory factor (*Mif*), interleukin 13 (*Il13*); measured by PCR) (Jia et al., 2016). Notably, in the latter study, the beneficial effects of RGFP966 on the RotaRod task was observed in female HD mice only, suggesting potential differences in HDACi’s effects depending on sex. While a sex comparison was not within the scope of our study, it would be important to test RGFP109, which showed therapeutic effects in male R6/1 mice in this current study, also on female R6/1s in future. In a mouse model of Friedreich ataxia prolonged RGFP109 treatment led to modest improvements in motor coordination performance and locomotor activity (Sandi et al., 2011). It has been shown that improvement on the RotaRod mainly requires a change in the motor strategy to master the task rather than an enhancement in general locomotor activity (Buitrago et al., 2004). Accordingly, we did not observe improvement in general locomotor activity in the open field test upon RGFP109 treatment in R6/1 animals.

To investigate the potential molecular correlates of our behavioral findings, we carried out a genomewide analysis of the transcriptional changes, which was not done before in the afore-mentioned studies, in the striata of vehicle- and drug-treated R6/1 and wild type mice. Our RNA-seq data showed the altered expression of a large number of genes in the striatum of R6/1 mouse, which included also those that are well-known to be dysregulated in HD, e.g. *Drd1, Drd2, Penk* and *Adora2a* (Seredenina and Luthi-Carter, 2012). RGFP109 administration showed, in part, repair effects on the global aberrant gene expression changes in the striatum of R6/1 mice, affecting 43 individual genes significantly. Some of these genes were the thyrotropin-releasing hormone gene *Trh*, the neuropeptide-like molecule gene *Nxph3*, the nicotinic acetylcholine receptor gene *Chrna4*, and transcriptional regulator genes such as *Neurod2, Neurod6, Nr4a2* and *Satb2*, whose change in expression may likely result in alterations in the expression of their target genes. Further, *Arhgap15* (Rho GTPase activating protein 15) and *Gsto2* (Glutathione S-Transferase Omega 2), genes whose expression were completely rescued in a previous report by the HDAC1 and HDAC3 selective inhibitor HDACi 4b in R6/2s (Thomas et al., 2008), were also restored in our study upon RGFP109 treatment in R6/1s, supporting a similar mode of action of these selective HDAC inhibitors across different models of HD. Of note, loss of *Arhgap15* gene, a member of the Rac signaling pathway, was shown to cause decreased synaptic density and cognitive deficits in mouse and its mutations were identified in association with neurological and cognitive deficits in patients with Intellectual Disability (Zamboni et al., 2018), revealing the rescue of its expression by HDACi treatment as potentially therapeutically relevant in HD. Beyond these single gene expression changes, we found that a number of gene sets associated with neurophysiological functions relevant for HD pathology were changed upon treatment and these changes could be linked to the observed phenotypic changes. Also, it should be taken into consideration that gene expression changes restricted to certain cell types in the striatum may have been diluted in RNA-seq measurements that use total striatal tissue preparations as in the present study.

In an animal study of Friedreich Ataxia it was shown that long-term RGFP109 treatment (100 mg/kg over a period of five months with five injections per week) increased especially local H3K9ac and H4K5ac levels directly at the frataxin gene accompanied by higher frataxin gene expression levels in mouse brain, whereas global H3 and H4 acetylation patterns did not significantly increase by the drug (Sandi et al., 2011). Next to the spatial changes of specific histone acetylation patterns, temporal differences of global and local histone acetylation have been observed in another Friedreich Ataxia mouse study using a single RGFP109 administration (with 150 mg/kg) (Rai et al., 2010). In this study, global H3 acetylation in the brain increased to a maximum level at 4 hours after injection and totally disappeared at 24 hours, whereas H4K5ac and H3K14ac at the frataxin gene were shown to increase between 12 and 24 hours (Rai et al., 2010). In contrast to the aforementioned studies in mouse models of Friedreich Ataxia, we did not observe increased global H3K27 acetylation by RGFP109 treatment measured by Western blotting of total striatal extracts. Supporting our findings of no change in global histone acetylation after RGFP109 treatment, a previous report showed histone acetylation changes only at specific promoters using a similar HDAC inhibitor, HDACi 4b, in the R6/2 mice (Thomas et al., 2008).

Aiming at examining different key features of HD, on which RGFP109 treatment may have an effect, we performed MRI measurements to study the structural brain changes in HD mice. By registering the Allen brain atlas to MR images, we provided a complete list of regional and sub-regional volumetric changes of the brains of R6/1 animals, which by far expands the set of regions so far shown to be changed in this HD model. Using longitudinal *in vivo* MRI, a previous study detected reduction in both global brain volume as well as brain sub-regional volumes when corrected for global volume change in R6/1 mice over time, showing ubiquitous shrinkage of the striatum and the somatosensory cortices (Rattray et al., 2013), comparable with what we detected with our approach here. In contrast, increases in regional and sub-regional volumes when corrected for whole brain volume should be viewed cautiously, as they indicate a less pronounced shrinkage of that brain area compared to the volume reduction of the whole brain, rather than an actual regional size increase. In pre-symptomatic and symptomatic HD patients progressive whole brain volume loss is evident (Tabrizi et al., 2012). Furthermore, in addition to the striatum, other brain regions known to be affected in HD patients, such as cerebral cortex, amygdala, hippocampus, and brainstem (Rosas et al., 2003), are also changed in R6/1 mice. Similar to the modest transcriptomic changes we observed upon RGFP109 treatment, volumes of specific brain regions and sub-regions showed only non-significant trends towards an alleviation upon HDAC1 and HDAC3 inhibition in R6/1 mice at this disease stage. Of note, these regions include some of the brain areas that were previously implicated in motor skill learning behavior in rodents, such as the amygdala, ectorhinal cortex, entorhinal cortex, pallidum, piriform area, and primary motor cortex (Badea et al., 2019; Scholz et al., 2015; Tamakoshi et al., 2014).

In summary, our study demonstrates that treatment with the HDAC inhibitor RGFP109 provides benefits on key phenotypic and molecular aspects of HD as well as potential effects on the brain atrophy observed in the R6/1 mice. Further studies should include also relevant immunohistochemical assessments of the brain for elucidating potential synaptic plasticity changes that may contribute to the observed phenotypic effects of HDAC inhibition in HD mice. Although the R6/1 line recapitulates several key features of HD, such as motor and cognitive deficits, transcriptional dysregulation, accumulation of mutant Htt aggregates and brain atrophy, establishing direct mechanistic links between molecular pathology and specific behavioral deficits has been a challenge so far (Rattray et al., 2013). Further work using an HD mouse model with slower progression, intervention earlier in the time course of pathology, a longer treatment regime with various doses or a combination of these factors may enhance the leveraging of the therapeutic potentials of selective HDAC inhibitors in HD. Nevertheless, we have demonstrated that HDAC1/3 inhibitor RGFP109 improved motor skill learning deficit and alleviated transcriptional dysregulation, characteristic disease features, in HD mice.

## Author contributions

K. H. and F.Y. designed research, K. H., S. M. and M. H. M. performed research, K. H., S. M., S. P. K., C.H., B.K. and P. B.-S. analyzed data, and K. H. and F. Y. wrote the manuscript.

The authors declare no conflict of interest.

## Acknowledgements

This work was funded by Deutsche Forschungsgemeinschaft (DFG), Exc 257 NeuroCure (to F.Y.). Funding to S.M., S.P.K., CH and P.B.S. was provided by the German Federal Ministry of Education and Research (BMBF, Center for Stroke Research Berlin 01EO1301), the BMBF under the ERA-NET NEURON scheme (01EW1811) and the DFG (research grant BO 4484/2-1 and HA5741/5-1 to CH). We thank the Department of Experimental Neurology, Charité – Universitätsmedizin Berlin for support with behavioral analyses, the Core Facility 7T Experimental MRIs, Charité – Universitätsmedizin Berlin for support with MRI measurements, the Scientific Genomics Platforms, Max-Delbrück-Center for Molecular Medicine (MDC) in the Helmholtz Association, Berlin for sequencing, SPARK Berlin for scientific advice, Heike Lerch and Cansin Belgin Peksen for excellent technical assistance, and Theresa Hartung for critically reading the manuscript.

## Materials and Methods

### Animals

Hemizygous R6/1 mice, expressing exon 1 of the human *HTT* gene with 116 CAG repeat expansions, were purchased from the Jackson Laboratory. R6/1 mice were maintained on a C57BL/6J background crossing male R6/1 with female C57BL/6J. Genotypes were determined by PCR analyses (R6/1-fwd: 5’-CCGCTCAGGTTCTGCTTTTA-3’; R6/1-rev: 5’-GGCTGAGGAAGCTGAGGAG-3’). Littermates were randomly divided into four groups: WT-vehicle (n=10), R6/1-vehicle (n=13), WT-RGFP109 (n=10), R6/1-RGF109 (n=12). All mice used in the present study were housed together in groups of maximal four animals; if possible, genotype and treatment were mixed in individual cages. Weight of the mice were monitored throughout the study. No animal was excluded due to excess weight loss (exclusion criterium: ≥25% weight loss compared to the weight at the start of the experiment). One animal (R6/1-vehicle) died during the course of the experiment (after completion of the MRI measurement and open field test, before performing Rotarod and elevated plus maze tests) due to unknown reasons. Animals were housed under pathogen free conditions with *ad libitum* access to food and water on a 12 h light/12 h dark cycle at constant temperature (22±2 °C) and humidity (55±10%). All animal experiments were approved by the local animal care committee of *Charité-Universitätsmedizin Berlin* and by the *Landesamt für Gesundheit und Soziales Berlin* (license number G0314/16) and conducted according to the institutional guidelines. All efforts were made to minimize all unnecessary suffering of animals. In line with our license for animal experiments and the 3Rs principles, for reducing the data variation and thereby keeping the required number of mice for the experiments to a minimum, only male mice were used in this study.

### RGFP109 treatment of mice

The HDAC1 and HDAC3 inhibitor RGFP109 (RG2833; CAS No. 1215493-56-3) was purchased from Selleckchem. RGFP109 was dissolved in dimethyl sulfoxide and diluted in 0.9% NaCl (1:2) directly before use (final concentration: 15 mg/ml). Mice were administered RGFP109 (30 mg/kg body weight) or an equal volume of vehicle solution by intraperitoneal injections five times a week for three weeks (total of 23 days) starting at the age of 11 to 14 weeks. All injections and behavioral tests were done around the same time of the day to avoid any biochemical and physiological changes over the experiments. Mice were deeply anesthetized using isoflurane and sacrificed by cervical dislocation 18 h after the final injection. Brains were removed and separated into right and left hemispheres. One hemisphere was flash frozen at −80 °C in methyl butane (Sigma-Aldrich) for immunohistochemical analysis, one was further dissected for RNA extraction and immunoblot analyses of striatum.

### Behavioral assessment

Motor coordination, balance as well as motor skill learning were assessed by accelerating RotaRod performance test. The open field test was used to examine general locomotor activity. Mouse emotional state, fearfulness, arousal, and anxiety were evaluated by the elevated plus maze test assessing exploration and motor activity in a new open environment. On the day of the individual tests, animals were moved to the test room 30 minutes prior to the start of the test to allow sufficient time to habituate. All data recorded, regardless of individual behavior, were used for the analyses.

At the end of the treatment course, mice were tested for forelimb and hindlimb clasping behavior by suspending each mouse by the tail, 20 cm above their home cage for up to 30 seconds (lack of any clasping behavior scored 0; rapid movement of forelimbs scored 0.5; forelimb clasping behavior towards the abdomen scored 1; forelimb and hindlimb clasping behavior towards the abdomen scored 2).

#### Accelerating RotaRod

One day before the actual test, mice were trained on the RotaRod (TSE Systems) three times at constant speed (4 rpm for 60 seconds) and two times with accelerating speed (4 to 40 rpm over a period of 5 minutes) with 30 minutes interval between the sessions. During the training mice were placed back on the rod if they have fallen down. On the following day data were recorded for four test trials using the TSE RotaRod software, which detects the latency to fall in seconds and the rod rotational speed at fall in rpm. The test trials were done by accelerating the rod from 4 to 40 rpm over a period of 5 minutes with at least 30 minutes interval between the sessions. The rod was cleaned between animal trials to remove any odors.

#### Open Field

The open field test was performed in a grey open-top 50 x 50 cm arena (height 40 cm) located in a sound-attenuated observation chamber. Mice were individually placed near the wall of the box and locomotor activity was measured as total distance travelled and average velocity over a period of 10 minutes. VideoMot2 Software, TSE Systems was used to track and record all animals (standard measuring mode based on center of gravity). Vertical activity (supported rearing) was manually assessed analyzing the recorded videos (experimenter was blinded for treatment and genotype). The field was thoroughly cleaned between the animals to avoid any odor.

#### Elevated plus maze

The elevated plus maze test was performed in a sound-attenuated observation chamber. The maze configuration consisted of five main regions: two open arms (29.5 x 5 cm), two closed arms (29.5 x 5 cm) and a middle area connecting the arms. The plus maze was raised 68 cm above the floor. Animals were placed in the center of the maze towards one of the closed arms. Movements were tracked (standard measuring mode based on center of gravity) and recorded for 5 minutes using VideoMot2 Software (TSE Systems). Number, duration, and path of visits to the open arms were calculated. The maze was rigorously cleaned between the animals to remove any scent clues left by the previous subject mouse.

### MRI measurements

MRI measurements were performed under 1-2% isoflurane anesthesia in a 70:30 nitrous oxide:oxygen mixture. Temperature was maintained through a circulating warm water system. Respiration rate was monitored during the measurements (Small Animal Instruments, Inc., Stony Brook, NY). T2-weighted images were acquired on a 7T MR scanner (PharmaScan 70/16 US; Bruker, Ettlingen, Germany) using a 20 mm quadrature volume resonator (Rapid Biomed). To cover the whole brain, a 2D T2-weighted RARE pulse sequence was used with 32 contiguous axial slices with 0.5 mm slice thickness and in-plane field of view of 25.6 x 25.6 mm. The imaging parameters were: matrix size 256×256, echo time spacing ΔTE=12 ms, repetition time TR / effective echo time TE = 4200 / 36 ms, bandwidth=46875 Hz, RARE factor 8, 4 averages, acquisition time 6:43 min.

The Allen brain atlas (Lein et al., 2007) was registered to individual MR images using the MATLAB toolbox ANTX (Koch et al., 2017) and the volume of each brain region was measured in mm^3^. For statistical comparison of brain region volume between groups a t-test (FDR, q<0.1) was applied. Only regions with a size of > 0.1mm^3^ (corresponding to 20 voxels) in at least one of the analyzed groups were included in the analysis. Both, absolute brain region volumes as well as brain region volumes normalized to whole brain volume were evaluated.

### Gene expression analysis by RNA-seq, RNA-seq data analysis, and qRT-PCR

#### RNA isolation

Flash-frozen tissues were homogenized in QIAzol Lysis Reagent (Qiagen) followed by RNA extraction using the miRNeasy Kit (Qiagen). Libraries for RNA-seq analysis were prepared using the TruSeq RNA kit from Illumina. Two pools with twelve libraries each (pool 1: WT-vehicle (n=3), R6/1-vehicle (n=3), WT-RGFP109 (n=3), R6/1-RGF109 (n=3); pool 2: WT-vehicle (n=2), R6/1-vehicle (n=3), WT-RGFP109 (n=3), R6/1-RGF109 (n=4)) were sequenced on a Illumina HiSeq4000 instrument (1×51bp).

#### RNA-seq mapping and analysis

Raw single-end reads of cDNA fragments were aligned to the mouse transcriptome (RefSeq, mm10) using the ‘RNA-Seq Alignment Workflow’ from BaseSpace, Illumina (version 1.1.0) with STAR aligner (version STAR_2.5.0b) for mapping and Cufflinks (version 2.2.1) for fragments per kilobase of exon per million fragments mapped (FPKM) estimation of reference genes. Differential gene expression analysis was performed using ‘Cufflinks Assembly & DE Workflow from BaseSpace, Illumina (version 2.1.0) with Cuffdiff 2 (Cufflinks, version 2.2.1). Differential expressed genes with FDR, q<0.1 were considered significant.

#### Gorilla analysis

Functional enrichment for gene ontology terms (biological process, molecular function, and cellular component) were calculated using the two unranked list approach (target and background lists) from Gorilla (Eden et al., 2009). As target gene list all genes significantly different in R6/1-vehicle / WT-vehicle comparison (FDR, q<0.1; log2FC>|0.5|) and affected by RGFP109-treatment in R6/1 mice by at least 20% (283) were used. All expressed genes, with FPKM>0.1 in at least one sample (18844) were used as background.

#### Gene set enrichment analysis

For gene enrichment analysis only genes with FPKM>0.1 in at least one of the groups analyzed were used. Pseudocounts of 0.05 FPKM were added to every gene to circumvent inflated fold changes at low expressed genes. Mouse gene symbols were converted into human gene symbols using gene IDs from BioMART-Ensemble (https://www.ensembl.org/biomart). Enriched genes sets were analyzed using the software from Broad Institute (GSEA, (Mootha et al., 2003; Subramanian et al., 2005) with MgSigDB gene set collections for gene ontology, KEGG, and REACTOME pathways). 1000 gene-set-wise permutations were performed to generate the null distributions. Genes were ranked in descending order based on log2 fold changes. Enrichment scores were calculated using the classic statistic (unweighted). Gene sets with FDR, q<0.1 were considered significant.

#### Quantitative PCR

cDNA synthesis was performed with M-MLV reverse transcriptase (Promega) and random hexanucleotides. mRNA expression levels were assessed by quantitative real-time PCR using SYBR Green dye-based PCR amplification (Thermo Fisher Scientific) and the QuantStudio 3 detection system (Applied biosystems). Primer sequences are listed in Table S1. mRNA expression levels were calculated relative to housekeeping gene *Actb* according to following equation: 2^(Ct(*Actb*)-Ct(target gene))^.

### Immunoblot analysis

Flash-frozen tissues were homogenized in RIPA buffer (50 mM Tris pH7.4, 150 mM NaCl, 1% Triton X-100, 0.1% SDS, 1% sodium deoxycholate and protease inhibitor cocktail (Thermo Scientific)), incubated for 2 h at 4 °C, sonicated and clarified (12,000 x g). Equal amounts of protein (20 μg) were subjected to SDS-PAGE and subsequently transferred to a nitrocellulose membrane. Membranes were blocked and then incubated with primary antibodies (anti-H3K27ac, abcam; anti-H3, Cell Signaling Technology; anti-β Actin, Cell Signaling Technology) at 4°C overnight. Staining with secondary HRP-conjugated antibody was performed at room temperature for 1 h. Detection of the membrane was carried out using SuperSignal™ West Dura Extended Duration Substrate (Thermo Fisher Scientific). Histone intensities were compared and normalized to beta-Actin intensities from the same blot using ImageJ (ImageJ software, NIH).

### Immunohistochemistry

The hemispheres were cut in coronal sections of 30 μm using a Leica CM1950 cryostat. Sections were dried on glass slides at room temperature before fixating in an Aceton:Methanol (1:1) solution for 10 min at −20 °C. After a short drying period, sections were rehydrated in PBS and blocked for 1 h in 10% goat-serum in PBS with 0.1% Triton-X 100. Huntington aggregates were stained at 4 °C over night (EM48, Millipore; 1:500, diluted in 10% goat-serum in PBS). Slides were washed in PBS and incubated with a secondary antibody (anti-mouse, Rhodamine Red-X- conjugated, Jackson ImmunoResearch; 1:1000, diluted in 10% goat-serum in PBS). After washing sections were stained with DAPI, washed, and mounted in Immumount.

Image acquisition was performed using a Nikon Spinning Disk Confocal CSU-X microscope equipped with a Nikon Plan Fluor 40x/1.3 DIC H N2 objective and Andor iXon3 EMCCD DU-888 Ultra camera. Images were acquired using Nikon NIS-Elements (version 5.10) imaging software.

For HTT-aggregate quantification, 8 spatial separated z-stack images (21 images with 500 nm z-steps) of one striatal section per animal were acquired and analysed. HTT aggregates were counted manually based on maximum intensity projections of each stack (ImageJ software, NIH). Experimenter was blinded for treatment and genotype. Individual nuclei within tissue sections were identified using the DAPI signal. Using ImageJ, sum slice projections were created for each z-stack image followed by background signal subtraction (rolling ball radius: 15 pixels). A binary image was created, and the watershed method was used to separate touching objects. Particles larger than 15 μm^2^ were defined as nuclei.

### Statistical Analysis

Behavioral, qPCR, immunohistochemistry, and immunoblot data were analyzed using the Graphpad Prism 6.0 and 8.0 software. The statistical tests included unpaired, two-tailed Student’s *t*-test,, twoway ANOVA followed by Tukey’s multiple comparisons test (for measuring the response to genotype and treatment), and mixed-effects analysis with two-stage linear step-up procedure of Benjamini, Krieger and Yekutieli as post-tests (for repeated measures data) as indicated in each figure legend. Statistical comparisons, values for n and p (unpaired, two-tailed Student’s *t*-test and two-way ANOVA) or q (mixed-effects analysis) are indicated in the figure legends. Differences with p<0.05 or q<0.05 were considered significant. Visualizations of RNA-seq and MRI data and their statistical results were done in R or with Graphpad Prism 6 software. We use the term ‘trend’ for referring to non-significant changes that are below the significance threshold in the respective experiments.

---

**Figure S1.**
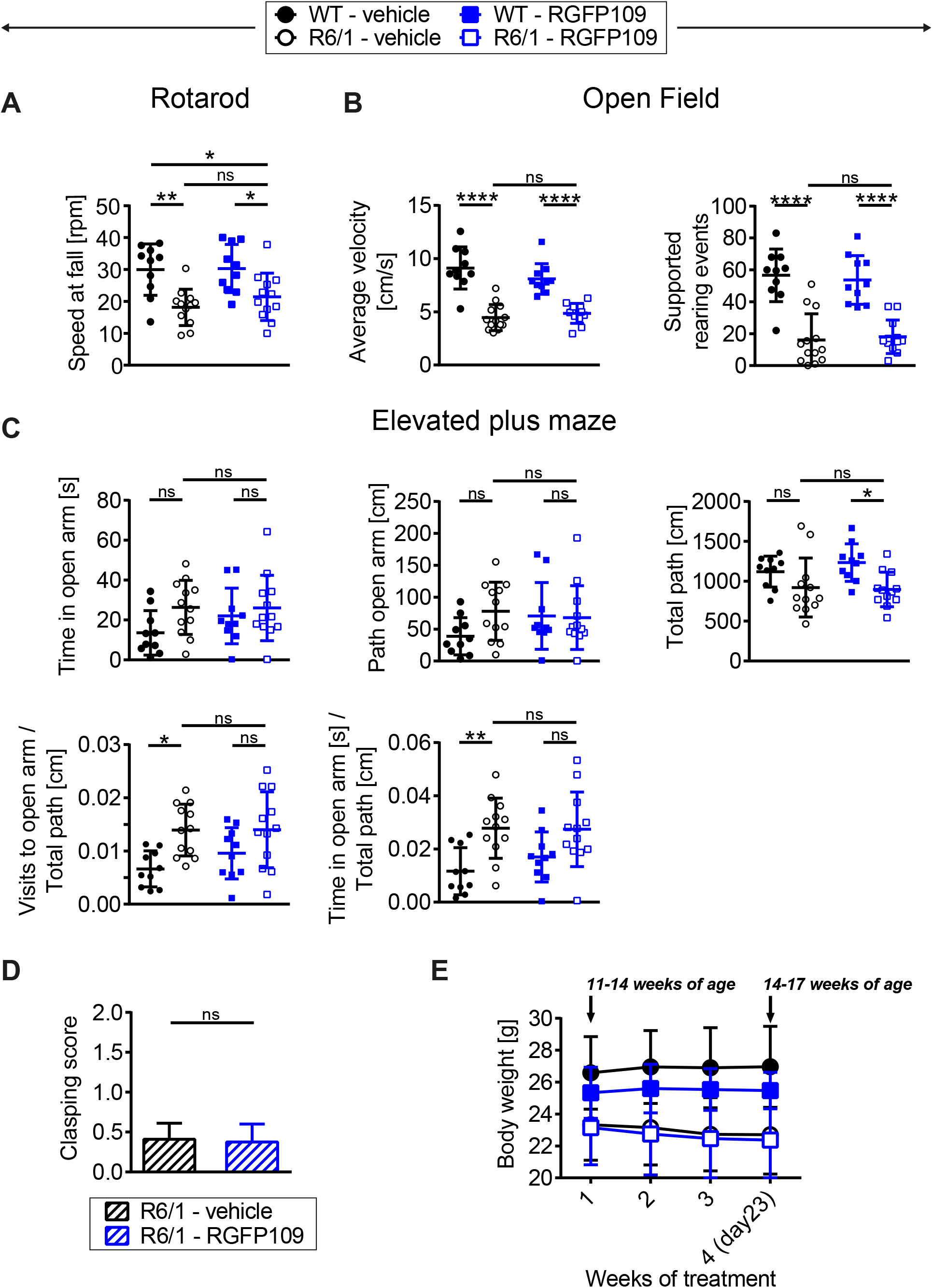
Behavioral and physical assessment of vehicle- and RGFP109-treated wild type and R6/1 mice. (A) Motor performance on a RotaRod: Speed at fall as mean of all trials. (B) Open field exploration evaluating general locomotor activity measured as average velocity and vertical activity (supported rearing). (C) Elevated plus maze test showing time in open arm, path in open arm, total path, and visits and time in open arm normalized to total path. (D) Clasping score of R6/1 mice measured at the end of the study. (E) Body weight over the course of the treatment. Data represent mean+SD; n=10 [WT-vehicle], n=12(13, Open Field) [R6/1-vehicle], n=10 [WT-RGFP109], n=12 [R6/1-RGF109]; *p ≤ 0.05, **p ≤ 0.01, ***p ≤ 0.001, ****p ≤ 0.0001, ns not significant by two-way ANOVA with Tukey’s multiple comparisons test (A-C) or by unpaired Student’s t-test (D).

**Figure S2.**
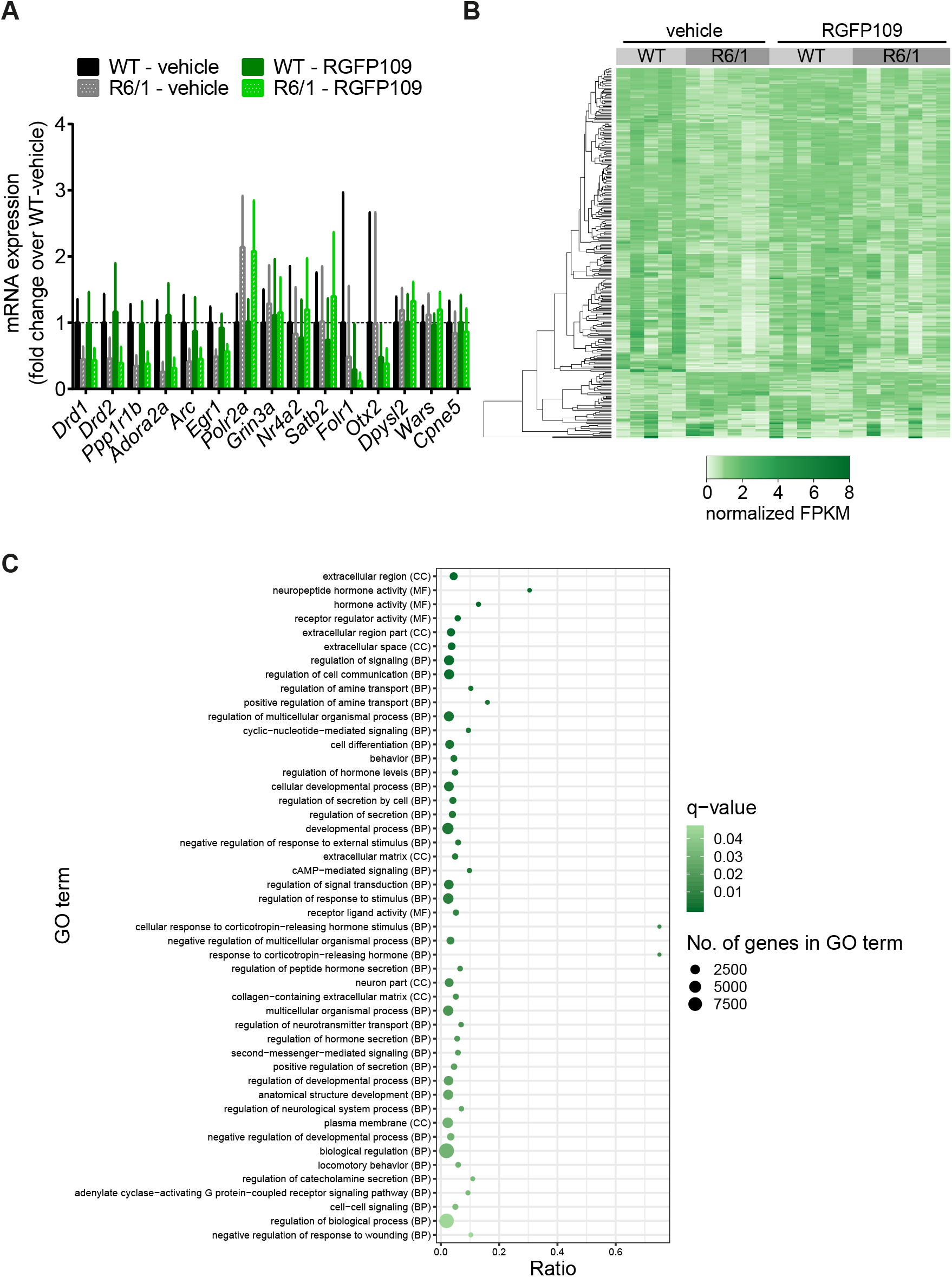
Effect of HDAC1 and HDAC3 inhibition by RGFP109 on gene expression in striatum of R6/1 mice. (A) Quantitative real-time PCR analysis of genes usually up- and downregulated in HD, motor-learning associated genes and those genes whose expression are found to be significantly altered in RNA-seq data by RGFP109 treatment of R6/1s in the striatum of vehicle- and RGFP109-treated wild type (WT) and R6/1 mice. Data represent mean+SD; n=10 [WT-vehicle], n=12 [R6/1-vehicle], n=10 [WT-RGFP109], n=12 [R6/1-RGF109]. (B) Heatmap showing differential expressed genes of R6/1 and WT comparison (FDR, q<0.1, log2 FC >|0.5|) which are changed towards WT levels by RGFP109 treatment in R6/1 mice by at least 20% (283). For better visualization FPKM values were mean-normalized. (C) Gene Ontology (GO) analysis of genes from (B). As background, all expressed genes with FPKM>0.1 in at least one sample were used (18844). Ratio describes the ratio between genes in the target gene list and the total number of input genes associated with a specific GO term. Enriched GO terms with FDR, q<0.05 are plotted; BP – biological process, CC – cellular component, MF – molecular function. n=5 [WT-vehicle], n=6 [R6/1-vehicle], n=6 [WT-RFGP109], n=7 [R6/1-RGFP109].

**Figure S3.**
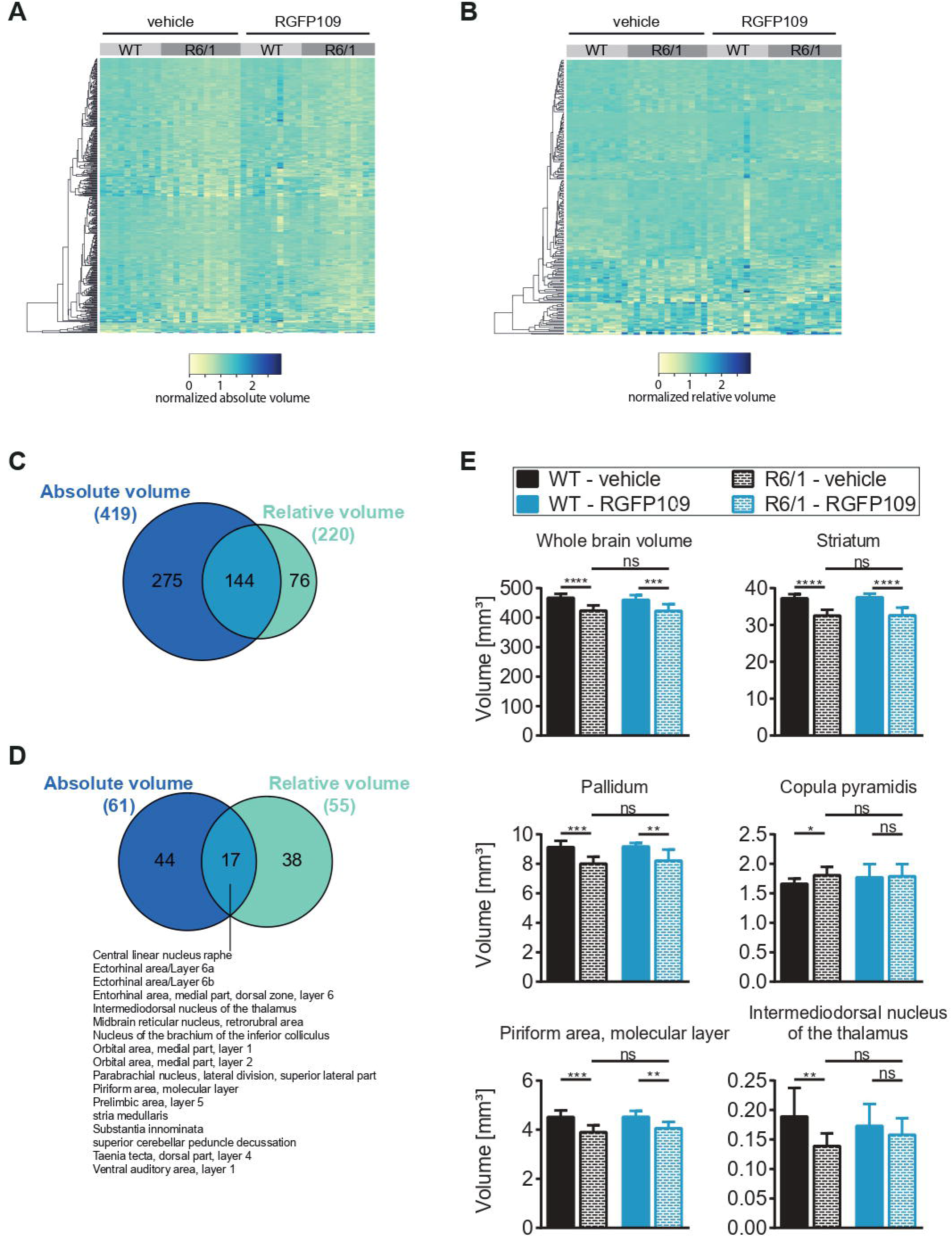
Influence of RGFP109 treatment on brain substructures in R6/1s measured by volumetric MRI. (A, B) Heatmaps illustrating absolute (A) or relative (B) volumes of regions with significant volume difference in WT-vehicle and R6/1-vehicle comparison (419 regions for absolute and 220 regions for relative volume). Volumes of every region were normalized to the mean volume of each region over all samples. (C) Venn diagram indicating shared and unique brain regions with significant volume change in R6/1 mice from absolute and relative volume analysis. (D) Venn diagram indicating shared and unique brain regions affected positively by the drug by at least 20% (not-significant) in R6/1 mice (red framed regions of Figure 3A and B) from absolute and relative volume analysis. (E) Absolute volumes of selected brain regions. Data represent mean+SD; n=10 [WT-vehicle], n=13 [R6/1-vehicle], n=10 [WT-RGFP109], n=12 [R6/1-RGFP109]; *q ≤ 0.05, **q ≤ 0.01, ***q ≤ 0.001, ****q ≤ 0.0001, ns not significant.

**Figure S4.**
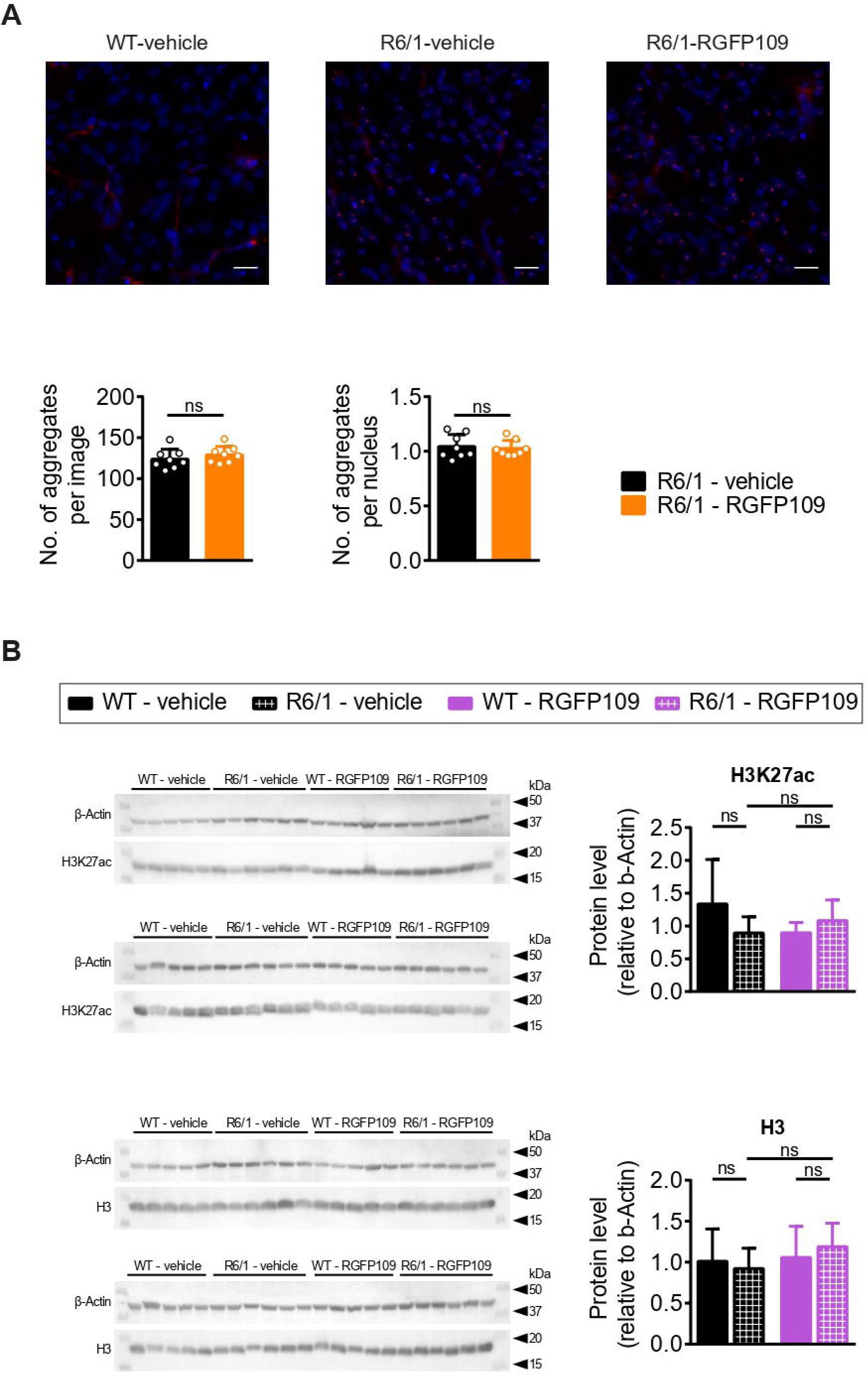

