## Supplementary material for "Selective inhibition of histone deacetylase 1 and 3 improves motor phenotype and alleviates transcriptional dysregulation in Huntington’s disease mice": Table S1

**Table S1. Primer sequences**

| Gene | Sequence (5' → 3') | |
|---|---|---|
| *Actb* | fw: | CACTGTCGAGTCGCGTCC |
| | rev: | TCATCCATGGCGAACTGGTG |
| *Adora2a* | fw: | GCCATCCCATTCGCCATCA |
| | rev: | GCAATAGCCAAGAGGCTGAAGA |
| *Arc* | fw: | TACCGTTAGCCCCTATGCCATC |
| | rev: | TGATATTGCTGAGCCTCAACTG |
| *Cpne5* | fw: | GTCTAACGGTGGTGTCCCAG |
| | rev: | TCCAGCTTGTTGGCACAGAA |
| *Dpysl2* | fw: | AAGCTGTGGTCACTGGGAAG |
| | rev: | GAAGACTTTGGCTGCGTTGG |
| *Drd1* | fw: | ATGGCTCCTAACACTTCTACCA |
| | rev: | GGGTATTCCCTAAGAGAGTGGAC |
| *Drd2* | fw: | AAGCGTCGGAAGCGGGTCAAC |
| | rev: | TCGGCGGGCAGCATCCATTCT |
| *Egr1* | fw: | TATGAGCACCTGACCACAGAGTCC |
| | rev: | CGAGTCGTTTGGCTGGGATAAC |
| *Folr1* | fw: | TTTACACGACCAGTGCAGCC |
| | rev: | TAGTTCCGCAGTGGTTCCAG |
| *Grin3a* | fw: | ACCAGTCAGAGGTTTCACAGAG |
| | rev: | GGTCCATCTTCTCCATCTGCTC |
| *Nr4a2* | fw: | GCAGAGAAGATCCCTGGCTT |
| | rev: | ACTGGGTTGGACCTGTATGC |
| *Otx2* | fw: | TCCAGGGTGCAGGTATGGTT |
| | rev: | AGCTCTTCTTCTTGGCAGGC |
| *Polr2a* | fw: | TGTGCAGGAAACATGACCGA |
| | rev: | GAAGCAGACACAGCGCAAAA |
| *Ppp1r1b* | fw: | CCAGAAACCCACTCTGTCCC |
| | rev: | GGCTTCAGCCAAAGCAAACA |
| *Satb2* | fw: | ACCGCACACAGGGATTATTGT |
| | rev: | CACTTCAGGCAGGTTGAGGA |
| *Wars* | fw: | AGCAGATCAAGAGCAAGGTCA |
| | rev: | TTCACAGTTGCCCCCAAACT |
